# Estimation of time-varying decision thresholds from the choice and reaction times with no assumption on the shape

**DOI:** 10.1101/090217

**Authors:** Yul Hyoung Ryul Kang

**Affiliations:** Department of Neurobiology, Columbia University

## Abstract

When a decision is made based on a series of samples of evidence, the threshold for decision may change over time. Here we propose a fast heuristic algorithm that estimates the time-varying thresholds without making an assumption on their shape. The algorithm gives an approximate best estimate of the time-varying thresholds for all time considered when all other parameters are fixed. The algorithm outperforms conventional methods that gradually adjust the threshold estimates during fitting, when fitting time is limited.

## Introduction

When a subject makes a binary choice based on a series of samples of scalar evidence, its speed and accuracy are typically modeled well by a diffusion process with two absorbing boundaries (Gold and Shadlen 2007; Bogacz et al. 2006). In these models, a decision is made when the cumulative sum of the evidence exceeds one of the two boundaries (or thresholds). It has been shown that when the quality of evidence is fixed (i.e., when there is only one signal to noise ratio), the number of accurate decisions per unit time can be maximized with thresholds that are constant over time (Wald 1973; Wald and Wolfowitz 1948). But the optimal form of the thresholds is different when signal to noise ratios vary between trials. There, trials with poor quality of evidence tend to take a long time to reach the threshold, and conversely, when a long time has passed and the threshold is not reached, it tends to be a trial with poor quality of evidence. Therefore, it is optimal to decrease the threshold over time, so as to move quickly onto the next decision, which is likely to be easier (Drugowitsch et al. 2012; Shadlen et al. 2006).

Although it has been reported that the normative solution for the optimal threshold over time can be found using the Bellman Equation (Drugowitsch et al. 2012), previous methods to estimate the shape of the thresholds had limitations. Many previous studies imposed a particular functional form (quadratic, Weibull, etc., see, e.g., Hawkins et al. 2015), and fit its parameters, but they may have missed the true shape of the thresholds if it cannot be expressed by the functional form chosen. At least one study used cosine basis functions (Drugowitsch et al. 2012), which can fit thresholds of any form given a sufficient number of basis functions. However, fitting the model is inefficient because it involves fitting many parameters using a gradient descent procedure, and it requires arbitrary hyperparameters, including the width of the cosine basis function.

Here we propose an efficient heuristic algorithm that does not require any parameter dedicated to thresholds. Although we defer rigorous justification of the algorithm to future works, we show that the algorithm can fit simulated data with no assumption on the shape of the thresholds, and do so significantly faster than previous methods with parametric thresholds.

## Methods

In the following text, we sometimes refer to the previous and the new approaches as parametric and nonparametric methods, following the way they specify the thresholds. Time and the level of accumulated evidence are discretized in steps of Δ*t* and Δ*y*. We ignore the discretization error as long as the error becomes arbitrarily small with smaller steps. We denote a random variable with an uppercase italic letter, and a scalar variable or constant with a lowercase italic letter. We denote a vector/matrix with a bold lowercase/uppercase letter, respectively (Table 1).

**Table 1.**
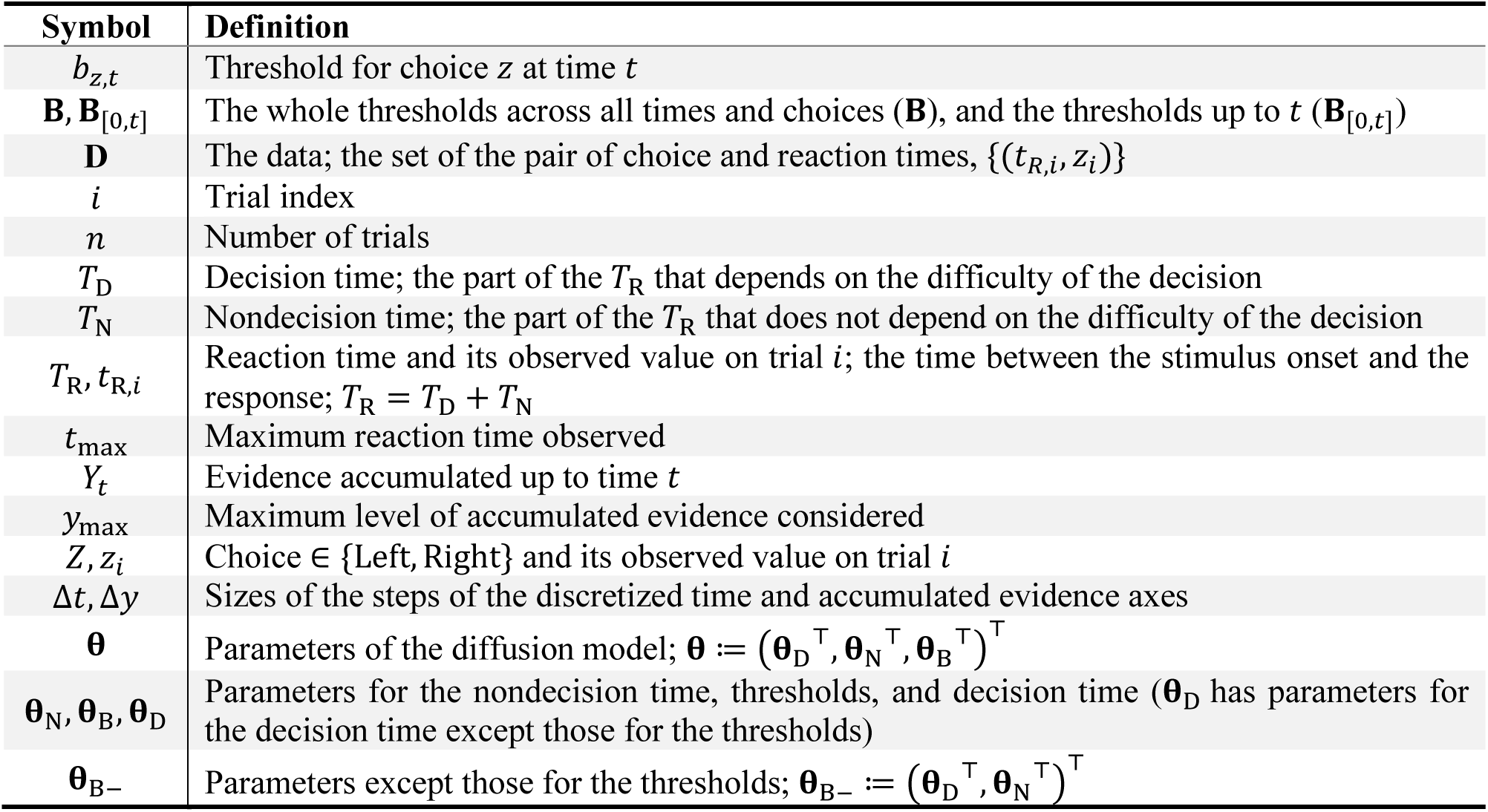
Important notations

| Symbol | Definition |
| --- | --- |
| $b_{z,t}$ | Threshold for choice $z$ at time $t$ |
| $\mathbf{B}, \mathbf{B}_{[0,t]}$ | The whole thresholds across all times and choices ($\mathbf{B}$), and the thresholds up to $t$ ($\mathbf{B}_{[0,t]}$) |
| $\mathbf{D}$ | The data; the set of the pair of choice and reaction times, $\{(t_{R,i}, z_i)\}$ |
| $i$ | Trial index |
| $n$ | Number of trials |
| $T_D$ | Decision time; the part of the $T_R$ that depends on the difficulty of the decision |
| $T_N$ | Nondecision time; the part of the $T_R$ that does not depend on the difficulty of the decision |
| $T_R, t_{R,i}$ | Reaction time and its observed value on trial $i$; the time between the stimulus onset and the response; $T_R = T_D + T_N$ |
| $t_{\max}$ | Maximum reaction time observed |
| $Y_t$ | Evidence accumulated up to time $t$ |
| $y_{\max}$ | Maximum level of accumulated evidence considered |
| $Z, z_i$ | Choice $\in \{\text{Left}, \text{Right}\}$ and its observed value on trial $i$ |
| $\Delta t, \Delta y$ | Sizes of the steps of the discretized time and accumulated evidence axes |
| $\boldsymbol{\theta}$ | Parameters of the diffusion model; $\boldsymbol{\theta} := (\boldsymbol{\theta}_D^\top, \boldsymbol{\theta}_N^\top, \boldsymbol{\theta}_B^\top)^\top$ |
| $\boldsymbol{\theta}_N, \boldsymbol{\theta}_B, \boldsymbol{\theta}_D$ | Parameters for the nondecision time, thresholds, and decision time ($\boldsymbol{\theta}_D$ has parameters for the decision time except those for the thresholds) |
| $\boldsymbol{\theta}_{B-}$ | Parameters except those for the thresholds; $\boldsymbol{\theta}_{B-} := (\boldsymbol{\theta}_D^\top, \boldsymbol{\theta}_N^\top)^\top$ |

### Algorithm 1. Fitting a diffusion model with parametric or nonparametric thresholds

// Equations are simplified to aid intuition. See text for full expressions.

Load the data **D** ← {(*t*_R,*i*_, *z*_*i*_)}

Initialize parameters for the diffusion and nondecision times, **θ**_D_ and **θ**_N_ // together called **θ**_B−_

**if** method=parametric:

Initialize parameters for the thresholds, **θ**_B_

**end**

**do**

Initialize P(*Y*_*t*=0_ = *y*, *T*_D_ > *t*) and *P*(*T*_D_ = 0, *Z* = *z*) // Eqs.10–11

**for** *t* **in** Δ*t* **to** *t*_max_ **do**

Compute the distribution of accumulated evidence assuming no thresholds at *t*,

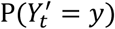, using P(*Y*_*t*−Δ*t*_ = *y*, *T*_D_ > *t* − Δ*t*) and **θ**_D_ // Eq. 12

**if** method=parametric:

Compute the threshold *b*_*t*_ using **θ**_B_ // Eq. 9

**else:** // method=nonparametric

Initialize the threshold *b*_*t*_

Compute the probability of outcomes **p**_*t*|**θ**_N__ that does not depend on *b*_*t*_,

using the data **D** and **θ**_N_ // Eqs. 19–21

**do**

Compute the probability of outcomes 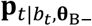 that does depend on *b*_*t*_,

Using 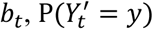, and **θ**_D_ // Eq. 22–24

Compute 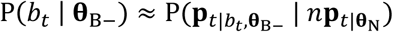 // Eq. 25

Adjust *b*_*t*_ to increase P(*b*_*t*_ | **θ**_B−_) // Eqs. 25–26

**until** P(*b*_*t*_ | **θ**_B−_) converges

**end**

Compute the probability of decision P(*T*_D_ = *t*, *Z* = *z*)

using 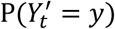 and *b*_*t*_ // Eqs. 14–15

Compute distribution of the accumulated evidence (with thresholds at *t*),

P(*Y*_*t*_ = *y*, *T*_D_ > *t*),

using 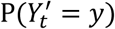 and *b*_*t*_ // Eq. 16

**end**

Compute the nondecision time distribution P(*T*_N_ = *t* | *Z* = *z*) using **θ**_N_ // Eq. 5

Compute the reaction time distribution P(*T*_R_ = *t*, *Z* = *z*)

using P(*T*_D_ = *t*, *Z* = *z*) and P(*T*_N_ = *t* | *Z* = *z*) // Eq. 3

Compute the cost, − ∑_*i*_ log P(*T*_R_ = *t*_R,*i*_, *Z* = *z*_*i*_ | **θ**), using P(*T*_R_ = *t*, *Z* = *z*) and **D** // Eq. 2

**if** method=parametric:

Adjust **θ**_D_, **θ**_N_ and **θ**_B_ to decrease the cost

**else:** // method=nonparametric

Adjust **θ**_D_ and **θ**_N_ to decrease the cost

**end**

**until** the cost converges

**return θ**_D_, **θ**_N_, and **B**

### Previous approach

We first review a previous, parametric approach to fitting the diffusion model. The key is that here the threshold heights are specified directly by parameters **θ**_B_, which will be introduced later in this section. In the next section, we will introduce a new, nonparametric method that proposes the threshold heights without any devoted parameter. The commonality and the difference between the two methods are summarized in Algorithm 1 and Figure 1.

**Figure 1.**
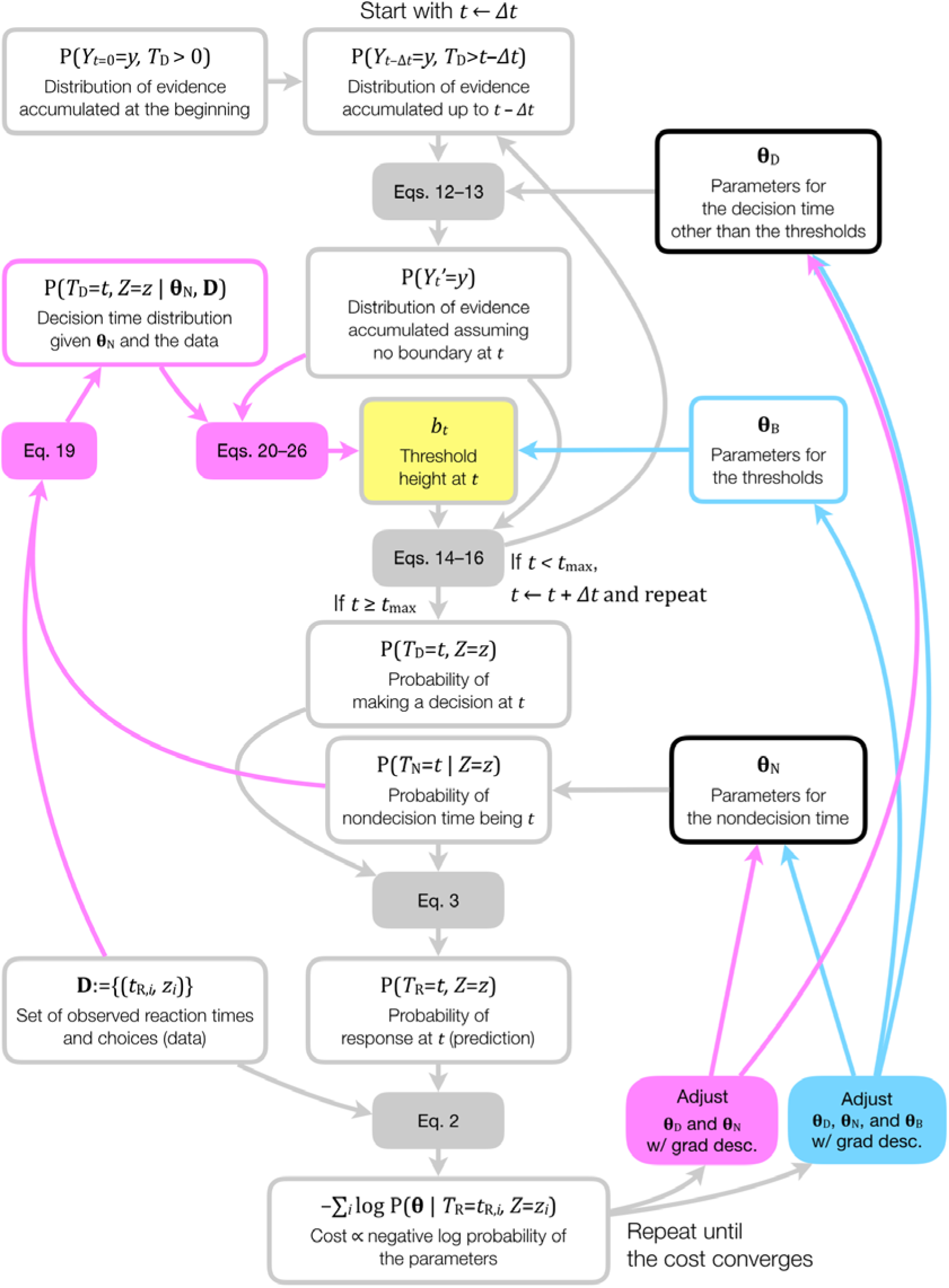
Flow of information. Expressions are simplified to aid intuition (see text for full expressions). Arrows denote dependence. Pink and blue colors denote quantities and dependence specific to the nonparametric and the parametric methods, respectively. Gray symbols are common to both. *b*_*t*_ is highlighted because it is the key quantity that the two methods calculate differently. Boxes denoting common parameters are colored black. “Grad desc.” indicates gradient descent.

For concreteness, we use an example of a direction discrimination task where on each trial the subject judges the direction of a motion stimulus to be left or rightward, although the mathematics holds for any 2-alternative forced choice experiment (Shadlen et al. 2006).

On each trial *i*, we observe a sample (*t*_R,*i*_, *Z*_*i*_) of the choice *Z* ∈ {Left, Right} and the reaction time *T*_R_ > 0, which is the time between the stimulus onset and the response. We decompose *T*_R_ into two parts, “decision time” (*T*_D_) that depends on the ambiguity of the stimulus (hence the difficulty of the decision) and “nondecision time” (*T*_N_) that consists of motor and sensory delays that are independent of the difficulty, such that

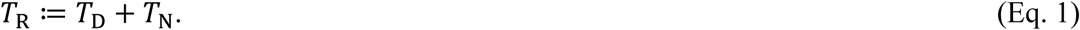

In previous studies, a vector of parameters **θ** was optimized to maximize the likelihood of the parameters given the data. Following the Bayes rule, it is the same as maximizing the log posterior probability of the data, or minimizing the negative log posterior, or “cost”:

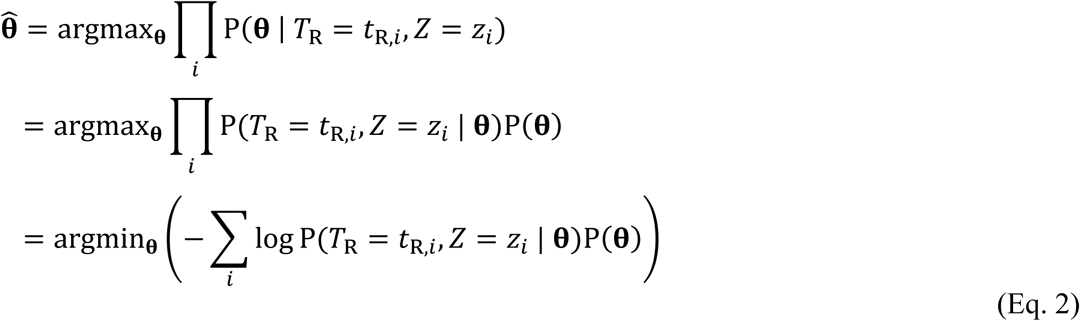

P(**θ**) is the prior, for which we use a relatively uninformative one that is flat within a range in this chapter (Table 2). We omit P(**θ**) in later equations for simplicity. P(*T*_R_ = *t*_R,*i*_, *Z* = *z*_*i*_ | **θ**) is the likelihood of observing the pair (*t*_R,*i*_, *z*_*i*_) on trial *i* given the proposed parameters. Note that the parameters **θ** include those that specify the thresholds, **θ**_B_: We can write **θ** ≔ (**θ**_D_^⊤^, **θ**_N_^⊤^, **θ**_B_^⊤^)^⊤^, where **θ**_N_ specifies the distribution of the nondecision time *T*_N_, and **θ**_D_ and **θ**_B_ jointly specifies the distribution of the decision time *T*_D_, where **θ**_D_ has the parameters that does not directly specify the thresholds.

**Table 2.** Free parameters

| Symbol | Definition | Initial value<br>(min–max) | # Used in models |  |  |
| --- | --- | --- | --- | --- | --- |
|  |  |  | Nonpara-<br>metric | Betacdf | Cosbasis |
| $\kappa$ | Gain of the drift | 5 (0.1–60) | 1 | 1 | 1 |
| $c_0$ | Bias of the drift | 0<br>(-0.256–0.256) | 1 | 1 | 1 |
| $\mu_{N,z}$ | Mean nondecision time for choice $z$ | 0.3 s<br>(0.15–0.75) | 2 | 2 | 2 |
| $v_{N,z}$ | Coefficient of variation of nondecision time for choice $z$ ($\sigma_{N,z}/\mu_{N,z}$) | 0.067<br>(0.05–0.95) | 2 | 2 | 2 |
| $b_0$ | Initial threshold height | 1 (0.01–3) | 0 | 1 | 0 |
| $t_\beta$ | Approximate time of the steepest collapse of the threshold | 0.5 s<br>(0.15–4.85) | 0 | 1 | 0 |
| $b_{\log}$ | Slope of the threshold | 1 (0–5) | 0 | 1 | 0 |
| $b_j$ | Coefficient for the $j$-th raised cosine basis function ($j \in [1, 7]$) | 0.5 (0–1.5) | 0 | 0 | 7 |
| $\beta_{b,1}, \beta_{b,2}$ | Parameter of the Betacdf that modulates the relative time | 0 (-0.5–0.5) | 0 | 0 | 2 |

The likelihood P(*T*_R_ = *t*_R,*i*_, *Z* = *z*_*i*_ | **θ**) in the last line of Eq. 2 is computed by convolving the predicted joint distribution of the choice and decision time P(*T*_D_ = *t*, *Z* = *z* | **θ**_D_, **θ**_B_) with the proposed nondecision time distribution P(*T*_N_ = *t* | *Z* = *z*, **θ**_N_), i.e.:

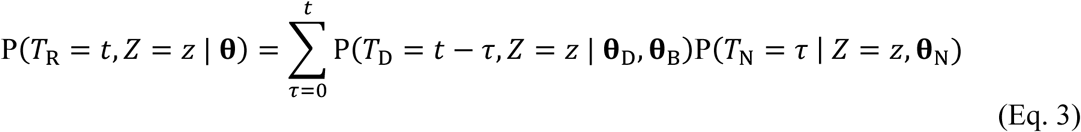

Because the dependence of the decision time on **θ**_B_ is only through the thresholds **B** that it specifies, we can write

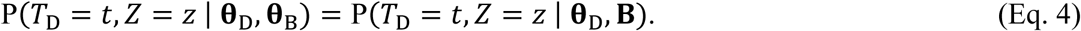

In this chapter, we use appropriately discretized gamma distributions for the nondecision time, parameterized in a way that **θ**_N_ consists of two pairs of numbers: the mean *μ*_N,*z*_ and the coefficient of variation *ν*_N,*z*_ (the ratio of the standard deviation and the mean), for each of the two choices *z*:

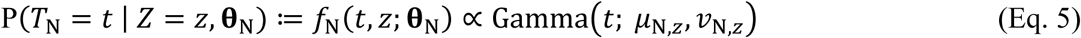

The likelihood of *T*_D,_ P(*T*_D_ = *t* − *τ*, *Z* = *z* | **θ**_D_, **B**), is computed by modeling *T*_D_ as the first crossing time of a one-dimensional diffusion process:

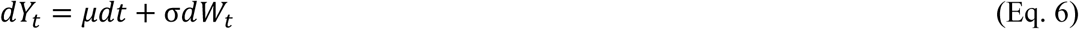

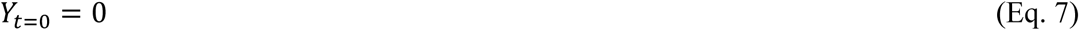

with two absorbing boundaries *b*_*z*=Left,*t*_ ≤ *b*_*z*=Right,*t*_ for the two choices, which in general vary over time. *μ* is the drift rate, *σ* is the diffusion coefficient, *Y*_*t*_ is evidence accumulated by time *t*, and *W*_*t*_ is the Wiener process. By convention, positive *Y* (and *μ*) supports the rightward choice, and negative leftward. Small absolute *μ* corresponds to a difficult decision. The decision time *T*_D_ is the first crossing time

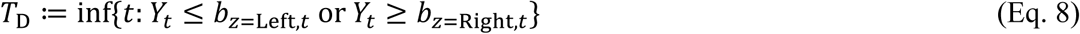

where inf{⋅} is the infimum. In previous approaches, the parameters **θ**_B_ specified the boundaries directly, i.e.,

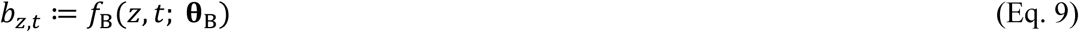

where *f*_B_(·) was, e.g., a beta cumulative distribution function (Kang et al. 2017) or a weighted sum of cosine basis functions (Drugowitsch et al. 2012). We will collectively denote the proposed boundaries across all times and choices **B**. (The key and the only difference of the new method is that **B** is not proposed using dedicated parameters **θ**_B_: it is instead inferred from the data using the rest of the parameters **θ**_N_ and **θ**_D_, as we will see in the next section.)

In the rest of this section, we will make concrete how *b*_*z*,*t*_ gives rise to the likelihood of *T*_D_. In this chapter, we use a rough yet simple approximation (for more accurate methods, see e.g., Smith 2000). We start with the initial condition, where at *t* = 0, no evidence is accumulated:

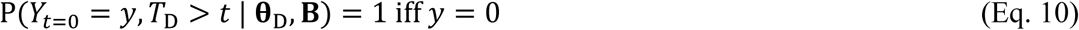

and no decision is made yet:

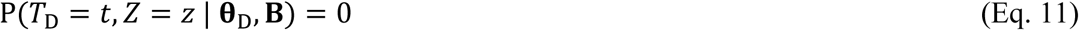

Then, for the next time step *t* ≥ *Δt*, we first compute the distribution of the accumulated evidence assuming no boundary (we denote such accumulated evidence as *Y*′ to distinguish it from the one *with* absorbing boundaries, *Y*):

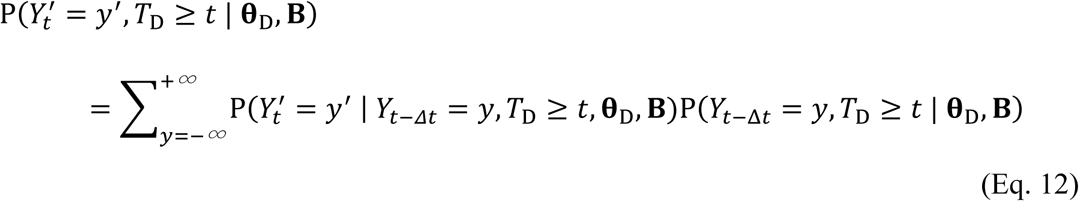

Here, note that P(*Y*_*t*−Δ*t*_ = *y*, *T*_D_ ≥ *t* | **θ**_D_, **B**)= P(*Y*_*t*−Δ*t*_ = *y*, *T*_D_ > *t* − Δ*t* | **θ**_D_, **B**) comes from Eq. 10 in the previous time step. The transition probability is parameterized by **θ**_D_ as a Gaussian distribution (and is independent of **B**, the latter because we are assuming no boundary for *Y*′):

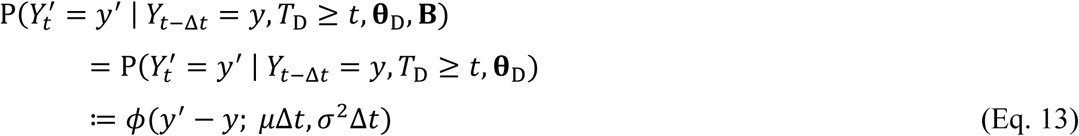

where *μ*Δ*t* is the mean, *σ*^2^*Δt* is the variance, *μ* is the drift rate, and *σ* is the diffusion coefficient. We fix *σ* to 1 without loss of generality (Palmer, Huk, and Shadlen 2005), leaving a single parameter, *μ*, in **θ**_D_. We then find the approximate probabilities of the rightward and leftward choice, at time *t* given a height of the threshold *b*_*z*,*t*_, as the sum of the probabilities outside the thresholds (for more accurate methods, see, e.g., Smith 2000):

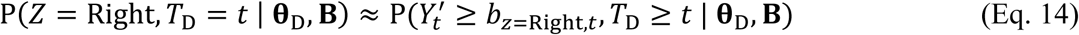

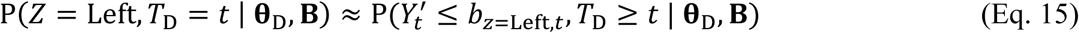

Then the probability of being undecided up to *t* with accumulated evidence *Y*_*t*_ *with* absorbing boundaries at *t* is set to that of the corresponding 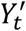 (the one without boundaries), except when 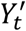 is outside the boundaries, in which case the probability is set to 0 (again, for more accurate methods, see, e.g., Smith 2000):

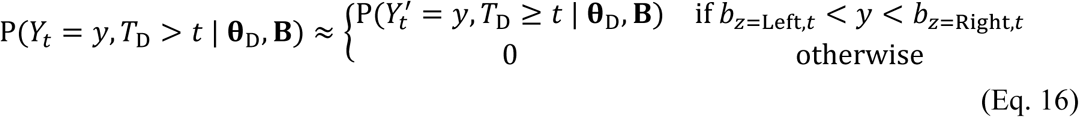

The steps in Eqs. 10–16 give the predicted probability of the decision time and the choice P(*T*_D_ = *t*, *Z* = *z* | **θ**_D_, **B**) for all *t* ≥ 0, which in turn gives P(*T*_R_ = *t*_R,*i*_, *Z* = *z*_*i*_ | **θ**) through Eq. 3, which is optimized as in Eq. 2. Now we are ready to see whxat is different in the new approach.

### Symmetric thresholds with one difficulty level

In this section, we introduce a new heuristic approach to propose the thresholds **B** without relying on **θ**_B_. We instead propose them by inferring them from the data, using the rest of the parameters **θ**_N_ and **θ**_D_, which we collectively denote as **θ**_B−_. Note that the proposal of the thresholds is the only difference of the new approach from the previous approaches explained in the previous section (Figure 1). In this section, we consider the case of symmetric thresholds, where *b*_*z*=Left,*t*_ = −*b*_*t*_ and *b*_*z*=Right,*t*_ = +*b*_*t*_, and consider the asymmetric case in a later section. We also consider only one drift rate (hence one difficulty level) for now, and generalize it to multiple difficulty levels in a later section.

To anticipate where we are getting at, we propose the threshold at each time step by maximizing the approximate probability of the threshold given the three possible outcomes (leftward, rightward, and no choice) at the time step, which we denote as P(*b*_*t*_ = *b* | **θ**_B−_) below (Eq. 17; Figure 2C). The challenge is that, because the time of the decision *T*_D_ (the first crossing time of the diffusion process, Eq. 8) is observable only through the reaction time *T*_R_ after a delay of the nondecision time *T*_N_ (Eq. 1), we do not know what the outcomes were at time *t*. So we infer the probabilities of the outcomes (leftward, rightward, and no choice) in two ways, one way that does not depend on *b*_*t*_, and the other way that does. The first way that does not depend on *b*_*t*_ is to use the parameters proposed for the nondecision time, and we put the resultant probabilities in a 3-vector, *p*_*t*_|**θ**_N_ (Figure 2A; Eqs. 19–21). The second way that does depend on *b*_*t*_ also depends on **θ**_D_ and thresholds proposed up to *t* − *Δt* (which we denote as **B**_[0,*t*−*Δt*]_), and we will put its results in another 3-vector, **p**_*t*|*b*,**θ**_B−__ (Figure 2B; Eqs. 22–24). Then

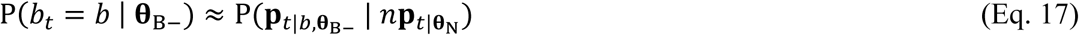

where *n* is the total number of trials (Figure 2C; see also Eq. 25). Then we set *b*_*t*_ to *b* that maximizes (the log of) this probability (Figure 2D; see also Eq. 26):

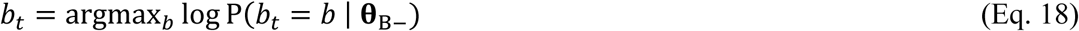

**Figure 2.**
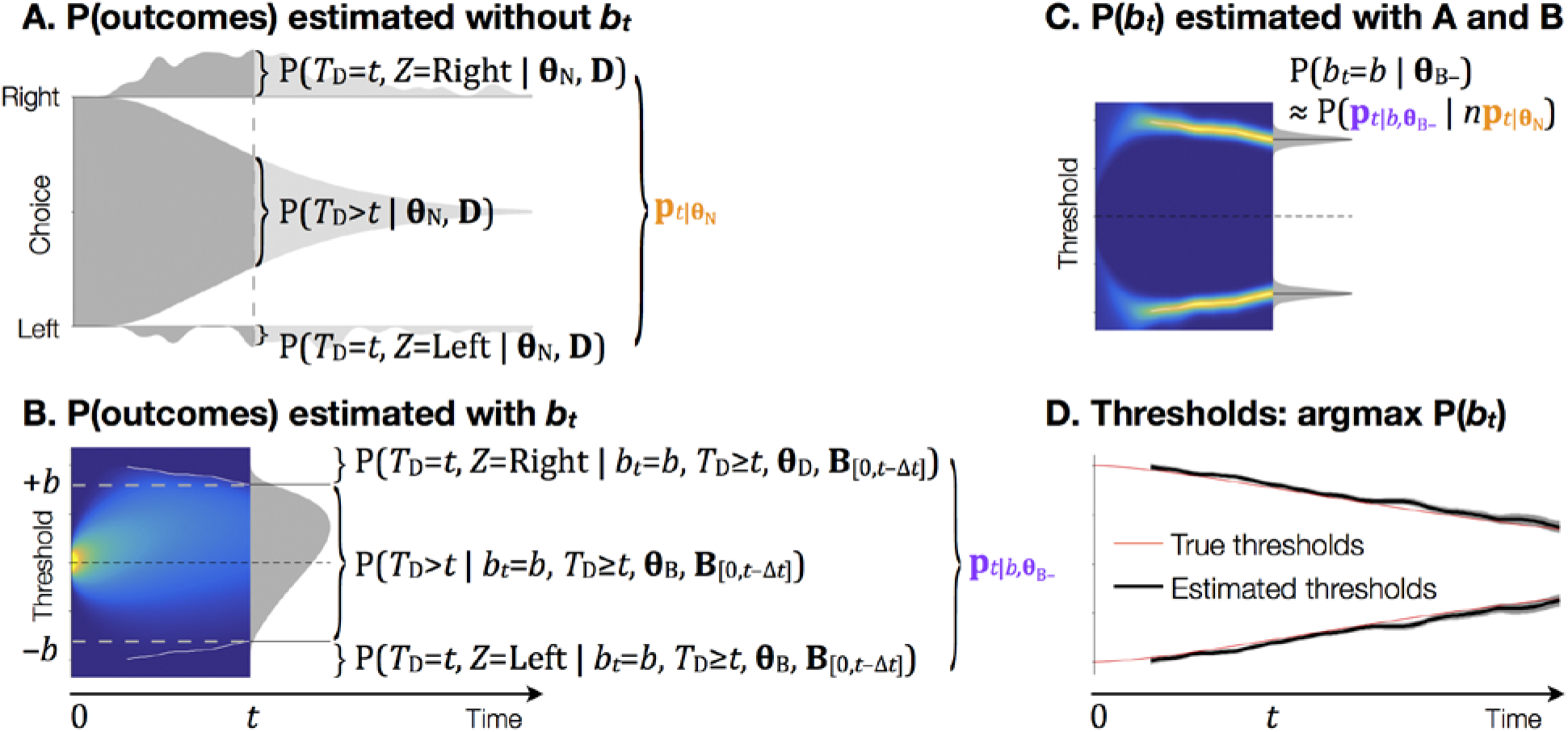
Estimation of the thresholds. **A.** Probability of the rightward, leftward, and no choice estimated *without b*_*t*_ (top, bottom, and middle shade and Figure 1, top pink frame), by convolving the observed distribution *T*_R_ with the proposed distribution of *T*_N_ flipped in time (Eqs. 19–21). Shades are lighter after *t* for visibility of the expressions. **p**_*t*|**θ**_N__ is a 3-vector of the three probabilities. B. Probability of the rightward, leftward, and no choice estimated *with b*_*t*_. Gray shade shows the distribution of the accumulated evidence assuming no threshold at *t*. If we impose the thresholds *b*_*t*_ = *b*, then the sums of the probabilities outside and within the thresholds approximate the probabilities of making a rightward or a leftward choice at *t* (top and bottom), and no choice (middle). **p**_*t*|*b*,**θ**_B−__ is a 3-vector that consists of the three probabilities, which sum to 1 (Eqs. 22–24). White solid lines are threshold heights estimated up to *t* − *Δt* (**B**_[0,*t*−*Δt*]_; displayed from the 5^th^-percentile of the estimated decision time distribution). Pseudocolor shows the distribution of the accumulated evidence for 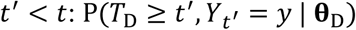. **C.** *Top half*. The likelihood of the threshold for the rightward choice being *b* at each time up to *t* (pseudocolor) and at *t* (gray shade; Eq. 25); *Bottom half.* The same for the leftward choice, which is a mirror image of the top, since we are assuming symmetric thresholds in this example. **D.** Thresholds estimated as the value maximizing P(*b*_*t*_ = *b* | **θ**_B−_) for each *t* (Eq. 26). Black solid lines are the maximum likelihood estimates, and shades are the standard errors (likely underestimated; see text after Eq. 26). The true thresholds used in the simulation are shown in red lines. The estimated thresholds are drawn between the 5^th^- and 95^th^-percentiles of the estimated decision time distribution.

Once we have the *b*_*t*_ to propose, the rest of the steps are the same as the previous approach (Figure 1, from *b*_*t*_ and below). To summarize, with the *b*_*t*_, we compute the decision time distribution for that time step, P(*Z* = *z*, *T*_D_ = *t* | **θ**_D_, **B**_[0,*t*]_), as in Eqs. 14–15 (we use **B**_[0,*t*]_ in the condition, since *b*_*t*_ became available in addition to the thresholds for the preceding time steps, **B**_[0,*t*−*Δt*]_.) We can repeat this procedure up to the maximum reaction time observed (*t*_max_) to get the entire decision time distribution P(*Z* = *z*, *T*_D_ = *t* | **θ**_D_, **B**) for all *t* ∈ [0, *t*_max_]. We in turn use it to compute the reaction time distribution P(*T*_R_ = *t*_R,*i*_, *Z* = *z*_*i*_ | **θ**_B−_) through Eq. 3, which is optimized as in Eq. 2 (with **θ** replaced with **θ**_B−_). So as mentioned earlier, the proposal of the thresholds is the only step that differs from the previous approaches: all other parameters, in **θ**_B−_ (**θ**_D_ and **θ**_N_), are optimized in the same way. We defer rigorous justification and analysis of the new method to future works, but the method gives reasonable fits fast, as we can see in Results.

In the rest of this section, we explain details of the new method outlined above. To compute the probabilities of the outcomes **p**_*t*|**θ**_N__ without relying on *b*_*t*_ (appeared at the end of Eq. 17 and Figure 2A), we start by computing the distribution of the decision times given the data **D** (a set of observed reaction times and choices {(*t*_R,*i*_, *z*_*i*_)}) and **θ**_N_. Because reaching the threshold for the choice *z* at time *t* would have happened in one of the trials where the choice *z* was made at a later time *t*_R_ ≥ *t*, with a probability of P(*T*_N_ = *t*_R_ − *t* | **θ**_N_), we can count all such trials weighted with the nondecision time probability to compute P(*T*_D_ = *t*, *Z* = *z* | **θ**_N_, **D**). In practice, we convolve the observed reaction time distribution with the nondecision time distribution reversed in time (not shown in the equation; the top pink frame in Figure 1 and Figure 2A):

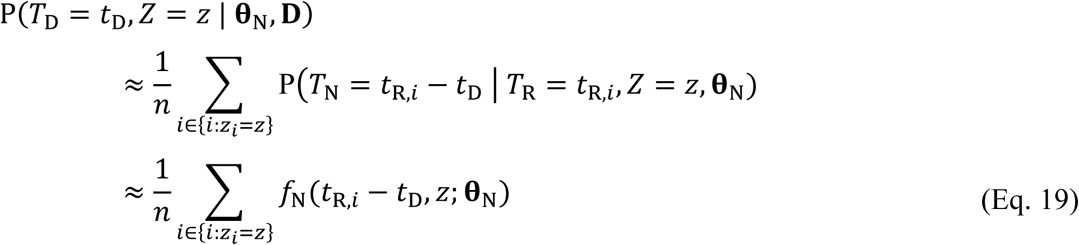

where *n* is the number of trials, {*i*: *z*_*i*_ = *z*} is the set of trial indices with choice *z*, **θ**_N_ is the vector of parameters regarding the nondecision time, and *f*_N_(*t*_N_, *z*; **θ**_N_) gives the probability of the nondecision time *t*_N_ given the choice *z* and parameters **θ**_N_, such that 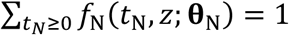 (Eq. 5). Obviously, *f*_N_(*t*_R,*i*_ − *t*_D_, *z*; **θ**_N_) = 0 ∀ *t*_R,*i*_ < *t*_D_.

Note that the decision time distribution computed this way is not guaranteed to maximize the probability of observing the data given **θ**_N_ (see Appendix). We still chose to use Eq. 19 because it is fast and simple, and we defer finding a better choice to future works. We confirmed that Eq. 19 nonetheless gives a better-than-chance approximation of the true simulated decision time distribution with a shuffle test (see Appendix and Figure 7).

With the decision time distribution estimated in Eq. 19, we also compute the probability that the decision is not made until *t* as (Figure 2A):

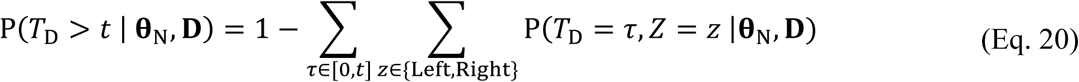

where square brackets denote a closed interval. Then we put the probability of leftward, rightward, and no choice into a 3-vector, **p**_*t*|**θ**_N__:

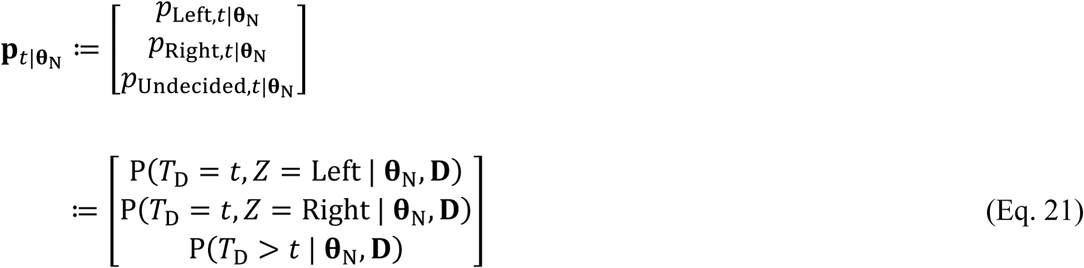

Note that the three elements do not sum to 1: they are the proportions of the trials out of all trials, and will be used to estimate the effective number of trials with each outcome at *t* (Eq. 25). The following equality may help: 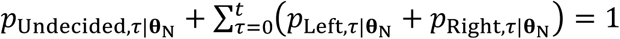 (see Eq. 20)

Now we compute the probabilities of outcomes that does depend on *b*_*t*_, **p**_*t*|*b*,**θ**_B−__ (appeared in Eq. 17 and Figure 2B). To do so, we first assume that we have already computed the probabilities of a decision and no decision up to *t* − *Δt*. For *t* = *Δt*, this is easy, since we know that no decision is made and no evidence is accumulated at *t* − *Δt* = 0, i.e., at the beginning (Eqs. 10–11). We can also assume *b*_*t*=0_ > 0, to avoid the trivial case where all decision is made immediately without any deliberation. For *t* > *Δt*, these probabilities depend on **θ**_D_, as well as the thresholds proposed up to *t* − *Δt*, **B**_[0,*t*−*Δt*]_ (which we assume we have, too.) Then we compute the probability of leftward and rightward choices given a certain value *b*_*t*_ = *b*, as we did in Eqs. 14–15 (Figure 2B): we first compute the distribution of accumulated evidence 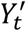 assuming no boundaries at *t* (Eq. 12), then impose the boundaries at ±*b* and sum the probabilities outside each boundary to get the probability of each choice. A slight difference is that we conditionalize with *T*_D_ ≥ *t* to make the probability of outcomes sum to 1, to make it work with the Dirichlet probability distribution function later (Eq. 25):

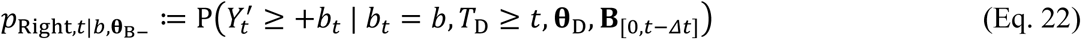

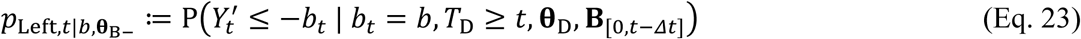

We also compute the probability that the decision is not made until *t* as the sum of the probabilities inside the two boundaries:

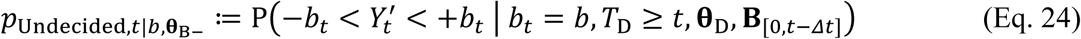

With the three probabilities in Eqs. 22–24, we construct a vector of length three, **p**_*t*|*b*,**θ**_B−__, whose elements sum to 1 (Figure 2B).

Once we have the two ingredients, **p**_*t*|**θ**_N__ and **p**_*t*|*b*,**θ**_B−__ for Eq. 17, we compute the approximate likelihood of the threshold height *b* heuristically, using the Dirichlet distribution (Figure 2C):

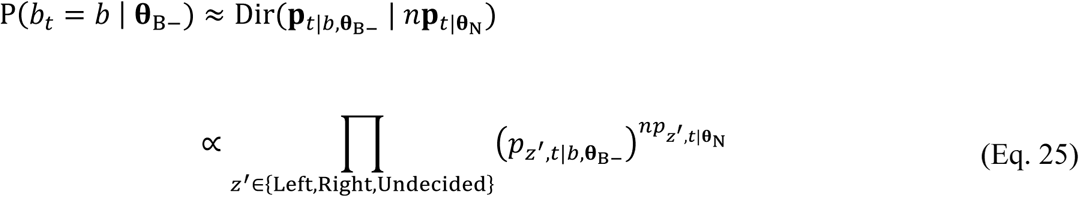

where 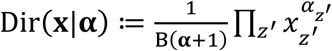 is the probability density function of the Dirichlet distribution where *z*′ indicates the category (in this case, *z*′ ∈ {Left, Right, Undecided}), *x*_*z*_′ is the probability of category *z*′ and **x** is a vector containing 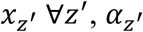 is the number of observed trials whose outcome is category *z*′ and **α** is a vector containing 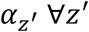, and B(**α** + 1) is the multivariate Beta function, which does not depend on *b* and hence is omitted in the last line of Eq. 25 (we define the elements of **α** to be smaller than usual by 1 for simplicity of the notation). The Dirichlet distribution gives the probability that, given the number of occurrences of categories in **α** sampled from a multinomial distribution, the probabilities of the categories are the elements of **x**. To make it concrete, note that the elements of **x** sum to 1, and the elements of **α** sum to 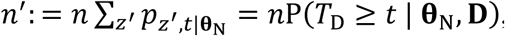, which is effectively the estimated number of trials that crossed the thresholds at or after *t*.

Then, as anticipated in Eq. 18, we set *b*_*t*_ as the value that maximizes its probability in Eq. 25 (Figure 2C and D):

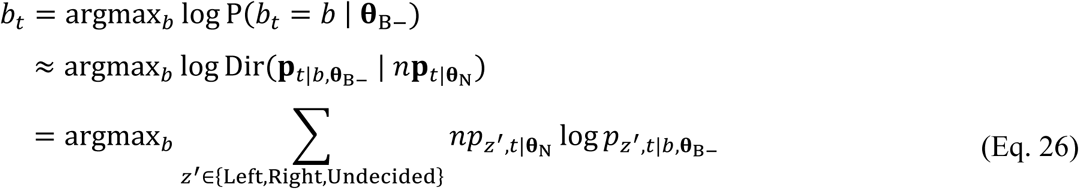

To see the rationale for Eqs. 25–26, imagine an extreme case where the nondecision time is a known constant 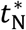, such that the threshold-crossing time on each trial is directly observable through the data, i.e., 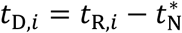 (i.e., there is no uncertainty about *n***p**_*t*|**θ**_N__). Further imagine that we have correct values of **θ**_D_ and of all thresholds up to *t* − *Δt*, **B**_[0,*t*−*Δt*]_, so that 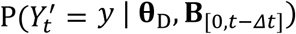 exactly matches the true distribution as well. Then, the probability that **p**_*t*|*b*,**θ**_B−__ gave to the number of “observations” *n***p**_*t*|**θ**_N__ (about which we have no uncertainty in this case), is exactly given by the Dirichlet distribution (to the extent that the approximation in Eqs. 14–15 holds, and except that we are ignoring the prior over *b*_*t*_). Since **p**_*t*|*b*,**θ**_B−__ in this case is a deterministic function of *b* (since we know **θ**_D_ and **B**_[0,*t*−*Δt*]_ with certainty), the probability of **p**_*t*|*b*,**θ**_B−__ equals the probability of *b*, making the approximation in Eq. 25 exact. Since the Dirichlet distribution is unimodal, given infinite amount of data *n* → *∞*, which makes the elements of **p**_*t*|**θ**_N__ exactly match the true value of P(*T*_D_ = *t*, *Z* = *z*) and P(*T*_D_ > *t*) under the generative model, Eq. 26 would give the true *b*_*t*_ that matches the generative model’s.

In practice, the preconditions are not met, at least not exactly, so we should use caution in interpreting P(*b*_*t*_ = *b* | **θ**_B−_) and *b*_*t*_ estimated from it. To begin with, neither **θ**_D_ nor **B**_[0,*t*−*Δt*]_ would match the true value initially in general, and **p**_*t*|**θ**_N__ estimated using Eqs. 19–21 is a rough and biased approximate when *T*_N_ has nonzero variance (see Appendix). Also, when the amount of data is finite, **B**_[0,*t*−*Δt*]_ would have variability, which is ignored here. Therefore, if we repeat the experiment with the same generative model, the variability of *b*_*t*_ estimated from Eq. 26 would be bigger than what P(*b*_*t*_ = *b* | **θ**_B−_) as in Eq. 25 suggests. This seem to be the case in our anecdotal observations, and we leave the full characterization of the confidence interval to future works. For now, we should keep in mind that the widths of the confidence intervals in Figures 2–4 are likely underestimated, which we noted in the legends.

Turning back to Eq. 26 with the caveats in mind, *b*_*t*_ for each time step can be found in *O*(log⌈*y*_max_/*dy*⌉) with a gradient ascent procedure with other parameters **θ**_B−_ fixed (*y*_max_ is the maximum level of accumulated evidence considered). It requires computing 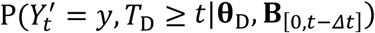 once per time step (as in the parametric methods), and we denote the time complexity for that with *O*(*s*), which is not smaller than *O*(⌈*y*_max_/*dy*⌉). Since *O*(log⌈*y*_max_/*dy*⌉) ≤ *O*(⌈*y*_max_/*dy*⌉), the new method does not increase the overall time complexity. (We measured and reported the execution time, as mentioned at the end of Section “Comparison of the performance with the parametric models” and corresponding Results). We do not provide a guarantee that the gradient ascent procedure will find the global maximum here, but in practice it worked reasonably, as we can see in Results.

As mentioned in the outset, we can repeat the above procedure, Eqs. 19–26, to obtain *b*_*t*_ for all time *t* ∈ [0, *t*_max_], and compute the decision time distribution P(*Z* = *z*, *T*_D_ = *t* | **θ**_D_, **B**) with Eqs. 10–16 and the reaction time distribution P(*T*_R_ = *t*_R,*i*_, *Z* = *z*_*i*_ | **θ**_B−_) with Eq. 3, and optimize it as in Eq. 2 (with **θ** replaced with **θ**_B−_). See Algorithm 1 for review.

### Symmetric thresholds with multiple difficulty levels

When there are multiple difficulty levels (in our example, different direction and noise levels of the motion stimulus), there are two additional steps we need to perform. First, we need to compute the drift rates of the diffusion process from the difficulty levels. For the motion direction discrimination task, the following linear transformation from the signed motion coherence *c* (positive when the motion is rightward and negative when it is leftward, with the absolute magnitude given by the probability of movement of each dot toward the signal direction; Shadlen and Newsome 2001) to the drift rate *μ*_*c*_ is known to work well (Palmer, Huk, and Shadlen 2005):

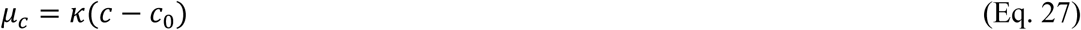

with two free parameters for **θ**_D_, the sensitivity *κ* and the bias *c*_0_.

Second, we need to pool results across difficulty levels. It can be done simply by summing the log likelihoods across conditions, and substituting it into Eq. 26:

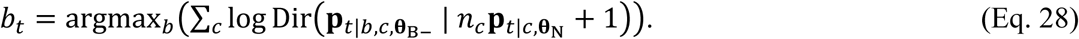

where *n*_*c*_ is the number of trials whose signed coherence is *c*, **p**_*t*|*b*,*c*,**θ**_B−__ is computed just like *p*_*t*|*b*,**θ**_B−__ with *μ* replaced with *μ*_*c*_ in Eq. 13, and **p**_*t*|*c*,**θ**_N__ is computed just like **p**_*t*|**θ**_N__ with trials whose signed coherence is *c*.

### Asymmetric thresholds

The method can be easily generalized to two asymmetric thresholds. Here, we calculate the likelihoods of the leftward, rightward and no decision as a function of both thresholds (cf. Eqs. 14–15 and 22–24):

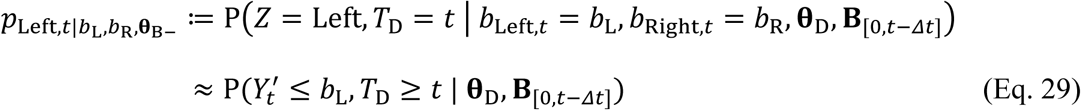

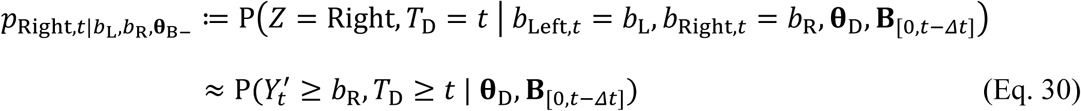

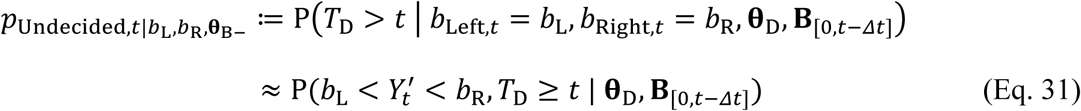

and with them, we construct a vector of length three, 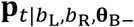. Then, as in Eq. 25, the likelihood of the thresholds is given by the Dirichlet distribution:

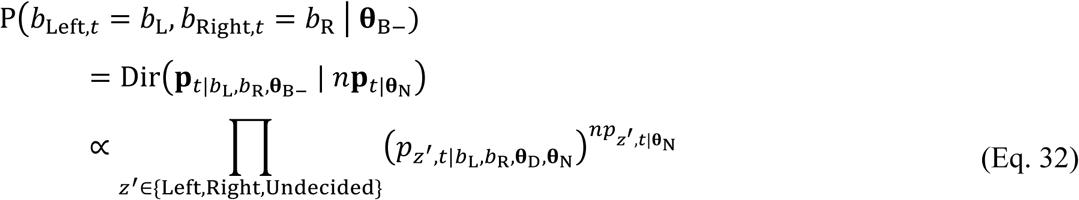

The threshold for *t* is set to the point with the maximum likelihood:

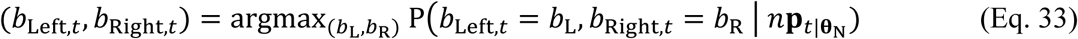

We can find the estimate using a gradient ascent procedure that optimizes (*b*_Left,*t*_, *b*_Right,*t*_) with other parameters fixed, with theoretical time complexity of *O*(⌈*t*_max_/*Δt*⌉ log *y*_max_), which is the same as the symmetric case. We defer characterizing the actual execution time to future works.

### Example fits

To evaluate the performance of the new method, we simulated the data by sampling the reaction time *T*_R_ and choice *Z* from P(*T*_R_, *Z* | **θ**) predicted by the diffusion model with known parameters (Eqs. 14–15), and fit them. First, as a sanity check, we tried giving the correct parameters **θ**_B−_ as the initial values (although not fixing them) and letting the new method find the thresholds (Figure 3, panels with the title “Original”, and Figure 4). We also tried making one parameter of the generative model different from the initial value of the free parameters and fitting them (Figure 3, other panels). See legends for generative parameters, and Table 2 for parameter settings used in fitting.

For absolute stimulus strengths |*c*|, we used 0, 0.032, 0.064, 0.128, 0.256, and 0.512, and generated positive and negative drift rates in a balanced way. For example, when we generated 1200 total trials, there were 100 trials for *c* = −0.512, 100 for *c* = +0.512, and so on, and there were 200 trials for *c* = 0 (because +0 = −0).

To parametrically generate collapsing thresholds, we used the same functional form as (Kang et al. 2017):

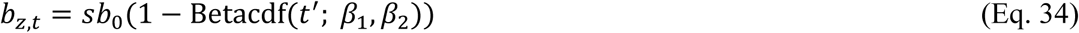

where 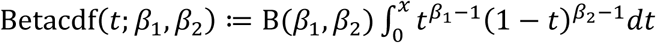 is the cumulative beta distribution function (where B(·) is the beta function, which gives a normalization factor), *t*′ ≔ *t*/*t*_max_ is the relative time, and the sign *s* is +1 for *z* = Right and −1 for *z* = Left (examples in Figure 3– Figure 4). In the following, we use *t*_β_ ≔ *t*_max_*β*_1_/(*β*_1_ + *β*_2_) and *b*_log_ ≔ log_10_(*β*_1_*β*_2_) instead of *β*_1_ and *β*_2_ to aid interpretation. Here, *t*_β_ is approximately the time of the steepest collapse (i.e., *t*_β_ ≈ argmax_*t*_ | *db*_*t*_/*dt* |), and *b*_log_ correlates with the slope of the collapse.

In generating the datasets with asymmetric thresholds (Figure 4), we used two sets of parameters for the thresholds, mentioned in the legend. When fitting the asymmetric thresholds data, we fixed *c*_0_ to 0, because a trend in both thresholds in the same direction (possible for asymmetric thresholds) is indistinguishable from a bias in the drift rate.

### Comparison of the performance with the parametric models

To systematically compare the performance of the new method (hereafter, the “nonparametric” method) with parametric methods (Figures 5 and 6), we generated the data with random settings of the threshold shapes and other parameters. Then we compared the cost (as in Eq. 2) between the nonparametric and the parametric methods on each step of gradient descent (“iteration”). For the comparison, we only used the symmetric thresholds, for which we could fit the model with a reasonable amount of data. It would be of interest to test the nonparametric method with asymmetric thresholds after improving it with extensions such as smoothness priors in future works.

We simulated the data with two models that differ in the shape of the thresholds used, which we call “betacdf” and “bump” thresholds and datasets. For the “betacdf” datasets, we used the same shape of the thresholds as in the previous section (Eq. 34 and Figure 5, left two columns). We used the smoothly and monotonically varying thresholds because their shape was similar to that of the commonly used parametric thresholds’, which we describe later in this section. We hypothesized that the parametric models would fit these datasets better than the nonparametric model. The parameters for the simulation were sampled independently and uniformly from a discrete set (denoted with curly braces) or from an open interval (denoted with parentheses) as follows: *κ* ∈ {5, 10, 20, 40}, *μ*_N_ ∈ (0.25, 0.4) s, *σ*_N_ ∈ {0.05, 0.1, 0.2} s (giving corresponding *ν*_*n*_ = *σ*_N_/*μ*_N_), *b*_0_ ∈ (0.5, 1.5), *t*_β_ ∈ (0.2, 1) s, *b*_log_ ∈ (0, 4).

For the “bump” datasets, we used thresholds with two step changes in the height in the opposite direction from each other (Figure 5, right two columns). We used the abruptly and non-monotonically varying thresholds because they did not match the shape of the thresholds of the common parametric models. We hypothesized that the nonparametric model would fit these datasets better than the parametric models. The initial height *b*_0_ was sampled uniformly from an open interval (0.5, 1.5). The two step changes were made at 0.5 and 1.0 s from the stimulus onset. When *b*_0_ < 1, the first step was upward and the second step was downward, and when *b*_0_ > 1, it was the opposite. The size of each step was sampled uniformly and independently from an open interval (0.2, 0.8). The parameters for the drift and nondecision times, *κ*, *μ*_N_, and *ν*_N_ were sampled in the same way as the betacdf datasets.

For each of the two generative models, we generated 180 random parameter settings (i.e., made 180 betacdf datasets and 180 cosbasis datasets), and for each parameter setting (i.e., for each dataset), we simulated 2400 total trials with the same set of stimulus strengths as in the previous section.

To fit the data, we used the nonparametric method as well as two parametric methods, and compared their performance (Figure 6). The nonparametric model was optimized as described in the previous sections until convergence.

The two parametric models differed only in terms of what functional form each used for the thresholds. One model used the beta cumulative distribution function, which is the same form as the betacdf datasets’, and we will call it the “betacdf” model. It had three free parameters for the thresholds, *b*_0_, *t*_β_, and *b*_log_, as described in the previous section. We will call the datasets generated with the betacdf thresholds as “betacdf datasets”, and fits made with the betacdf thresholds as “betacdf fits” to disambiguate them.

The other model used cosine basis functions, and we will call it the “cosbasis” model. For this model, we used the following function for the threshold (similar to Drugowitsch et al. 2012):

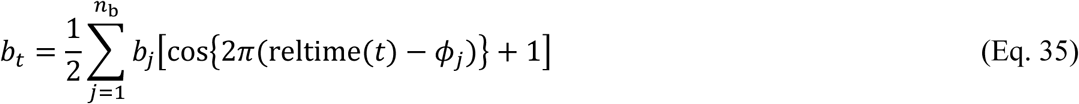

where cos(*x*) gives the cosine of *x* for – *π* ≤ *x* ≤ *π* and 0 otherwise. We used the number of the basis functions *n*_b_ ≔ 7, and spaced them 1/4 cycles apart from each other, by using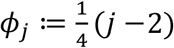. So it offered 7 free parameters for the threshold height, {*b*_*j*_}. All *b*_*j*_ were initially set to 0.5 (corresponding to *b*_*t*_ = 1 for all *t*). We also had two more parameters that define the relative time, reltime(*t*), which varies between 0 and 1 from the stimulus onset until the maximum time modeled (*t*_max_), following reltime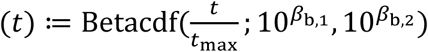, where Betacdf(·) is the cumulative beta distribution function as described in the previous section, and *β*_b,1_ and *β*_b,2_ are free parameters, both set to 0 at the beginning (which gives 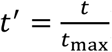). In total, the cosbasis model had 9 parameters for the thresholds.

All models, including the nonparametric model, had 6 additional free parameters regarding the drift (2) and the nondecision time (4). The starting point and the range of all free parameters are summarized in Table 2. Note that *c*_0_ was included as a free parameter in fitting following a common procedure for fitting the real experimental data, although we fixed it to 0 in simulating the data for simplicity. We also included *μ*_N,*z*_ and *σ*_N,*z*_ for each choice *z* ∈ {Left, Right}, although we set *μ*_N,*z*=Left_ = *μ*_N,*z*=Right_ and *σ*_N,*z*=Left_ = *σ*_N,*z*=Right_ in simulating the data, for the same reason.

To compare the performance as a function of the number of the iterations, we calculated the cost relative to the smallest cost at convergence across all three models within each simulated dataset (such that the minimum becomes 0 for each dataset). We then plotted the mean across the 180 datasets within each fitting method within each iteration and plotted it (Figure 6, first column). Because each method took different number of iterations until convergence, we used the cost at convergence for averages for iterations after convergence (e.g., if the cosbasis model converged on Iteration 30 with cost = 100 for Betacdf Dataset 15, we used 100 as the cosbasis model’s cost on all later [ > 30 ] iterations for Betacdf Dataset 15). We also plotted the percentage of simulations (among 180) that each parametric method had a lower cost than the nonparametric method on each iteration (Figure 6, second column).

While the nonparametric method offers the same time complexity as parametric methods, it requires extra time to search for the best threshold height per time step (Eq. 26). So we also computed the time per calculation of the cost for all three fitting methods, by averaging the time spent in 100 calculations after 3 warm-up calculations. Also, in addition to the number of iterations until convergence, we compared (1) the total execution time until convergence, and (2) the “pure” execution time devoted to the calculation of cost, by multiplying the number of iterations with the time spent per calculation of the cost for each fitting method.

## Results

We first ran a sanity check, by setting the initial values of all but 0–1 free parameter to the generative model’s and seeing if the new method can estimate the threshold. For symmetric thresholds, the method captured the shapes with as few as 600 total trials (Figure 3). In case of asymmetric thresholds, the method captured the qualitative difference between the two thresholds. Not surprisingly, the fit was worse compared to the symmetric case with the same number of trials (Figure 4, top row), but it improved with more trials (Figure 4, bottom row).

**Figure 3.**
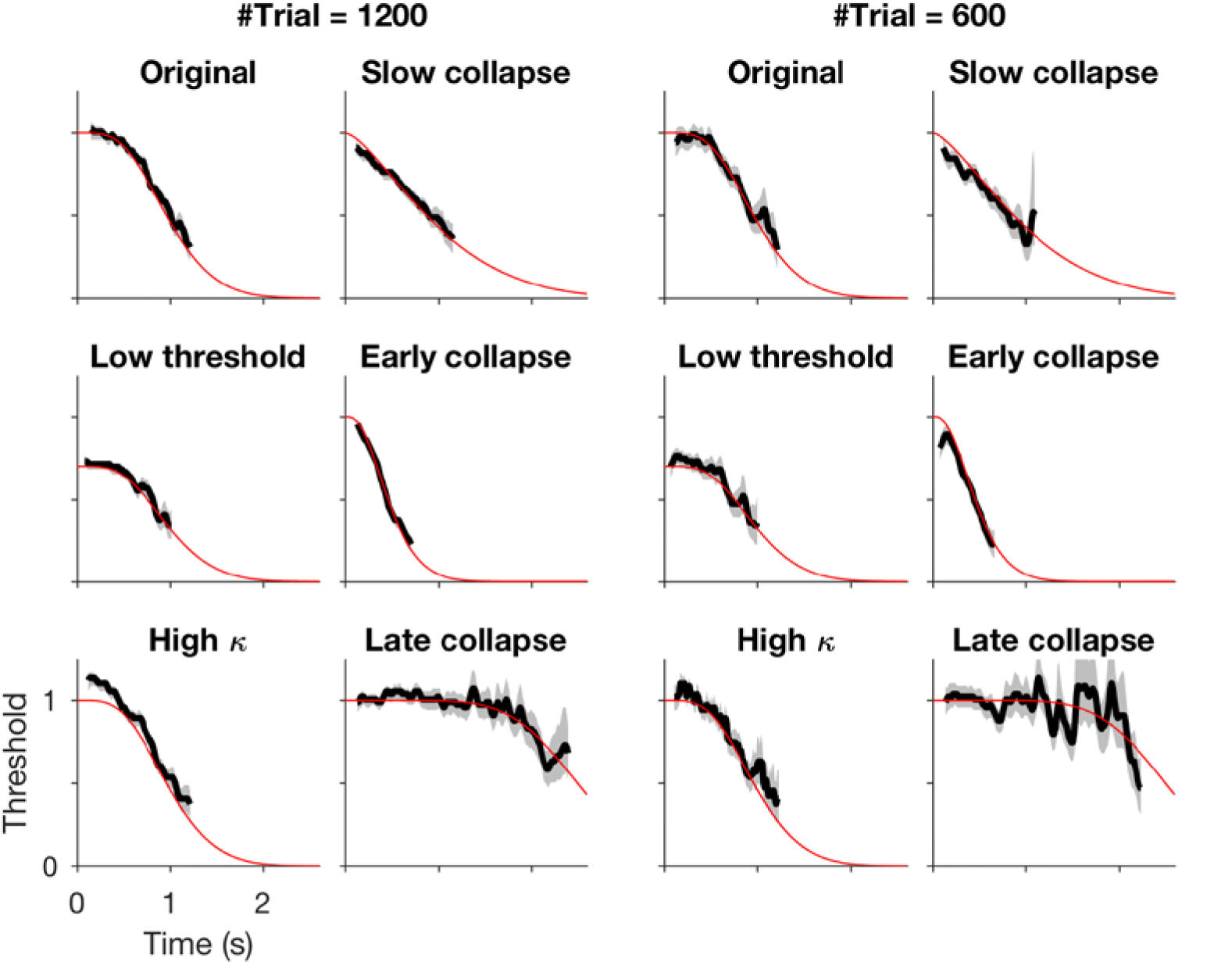
Comparison of the fit and the ground truth for symmetric thresholds. The black solid line is the maximum likelihood estimate, the shade is the 95% confidence interval based on P(*b*_*t*_ = *b* | **θ**_B−_) at each *t* (likely underestimated; see text after Eq. 26), and the red line is the true threshold. Estimates are plotted between 5 and 95 percentiles of the estimated decision times. Parameters used are: *κ* = 5 (or 10 for the “high *κ*”), *b*_log_ = 1 (or 0.5 for the “slow collapse”), *t*_*β*_ = 0.4 s (or 0.2 and 0.8 s for the “early collapse” and “late collapse”), and *b*_0_ = 1 (or 0.7 for the “low threshold”). All other parameters had the default values listed in Table 2. Left two columns are fit with the total of 1200 trials, right two columns with 600 trials.

**Figure 4.**
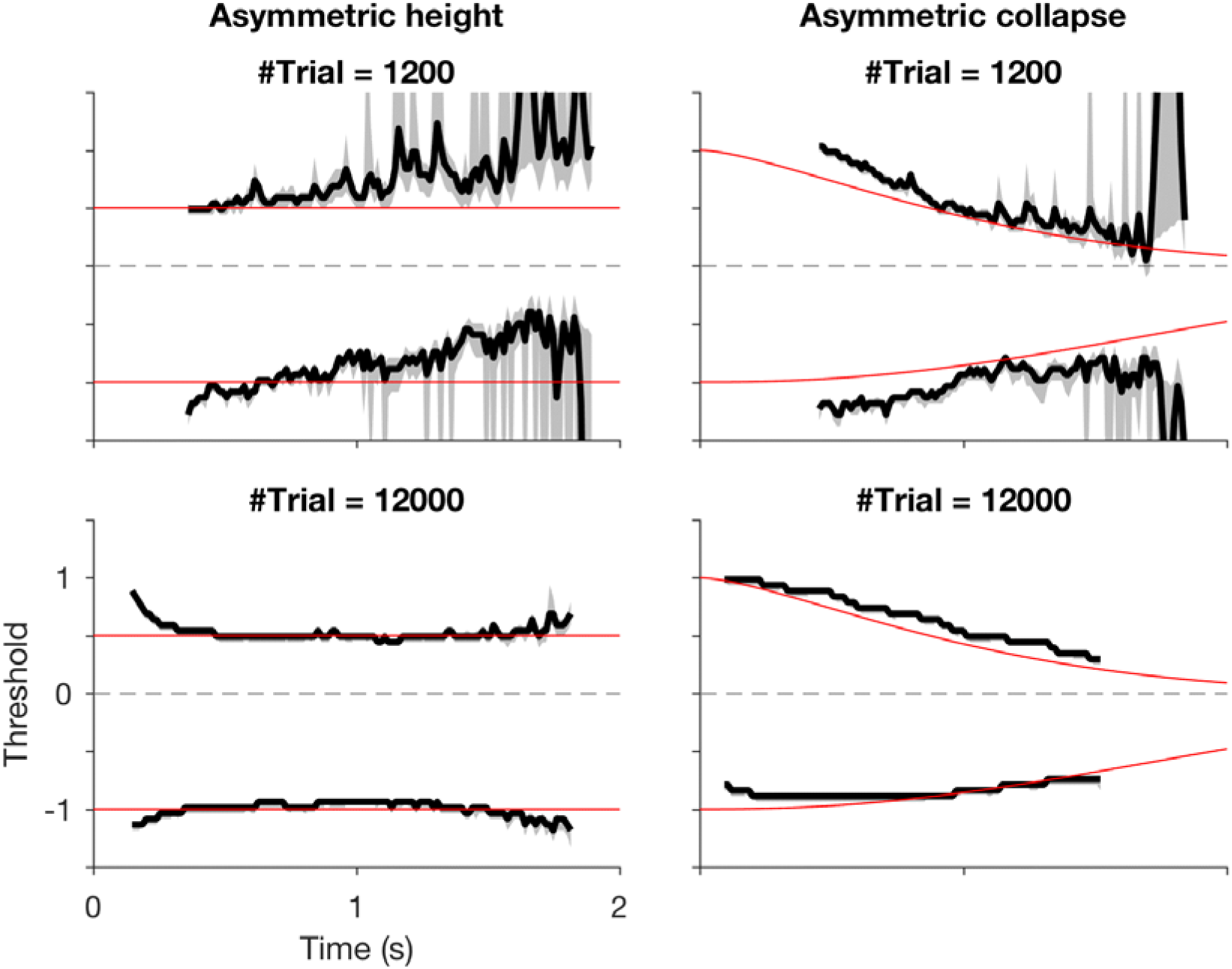
Comparison of the fit and the ground truth for asymmetric thresholds. The black solid line is the maximum likelihood estimate, the shade is the 95% confidence interval based on P(*b*_*z*,*t*_ = *b* | **θ**_B−_) for each *t* and *z* (likely underestimated; see text after Eq. 26), and the red lines are the true threshold. Estimates are drawn between 5^th^- and 95^th^-percentiles of the estimated decision times. The top row is fit with 1200 trials, bottom with 12000 trials. We used *κ* = 4 for both examples. For the “asymmetric height” examples, we used *b*_*z*=Right,*t*_ = 0.5 and *b*_*z*=Left,*t*_ = −1 for all *t*. For the “asymmetric collapse” examples, we used *b*_*z*=Right,*t*=0_ = *b*_*z*=Left,*t*=0_ = 1 and *t*_*β*,*z*=+1_ = 1 s and *t*_*β*,*z*=−1_ = 2 s, respectively. All other parameters had the default values listed in Table 2.

We then compared the performance of the nonparametric model with that of two parametric models, betacdf and cosbasis, with random parameter settings as a function of the number of iterations of the gradient descent (See Section “Comparison of the performance”). Visual examination of the fits revealed that the nonparametric fits capture the shape of the thresholds as early as the 2^nd^ iteration, for both the betacdf and bump datasets (Figure 5).

**Figure 5.**
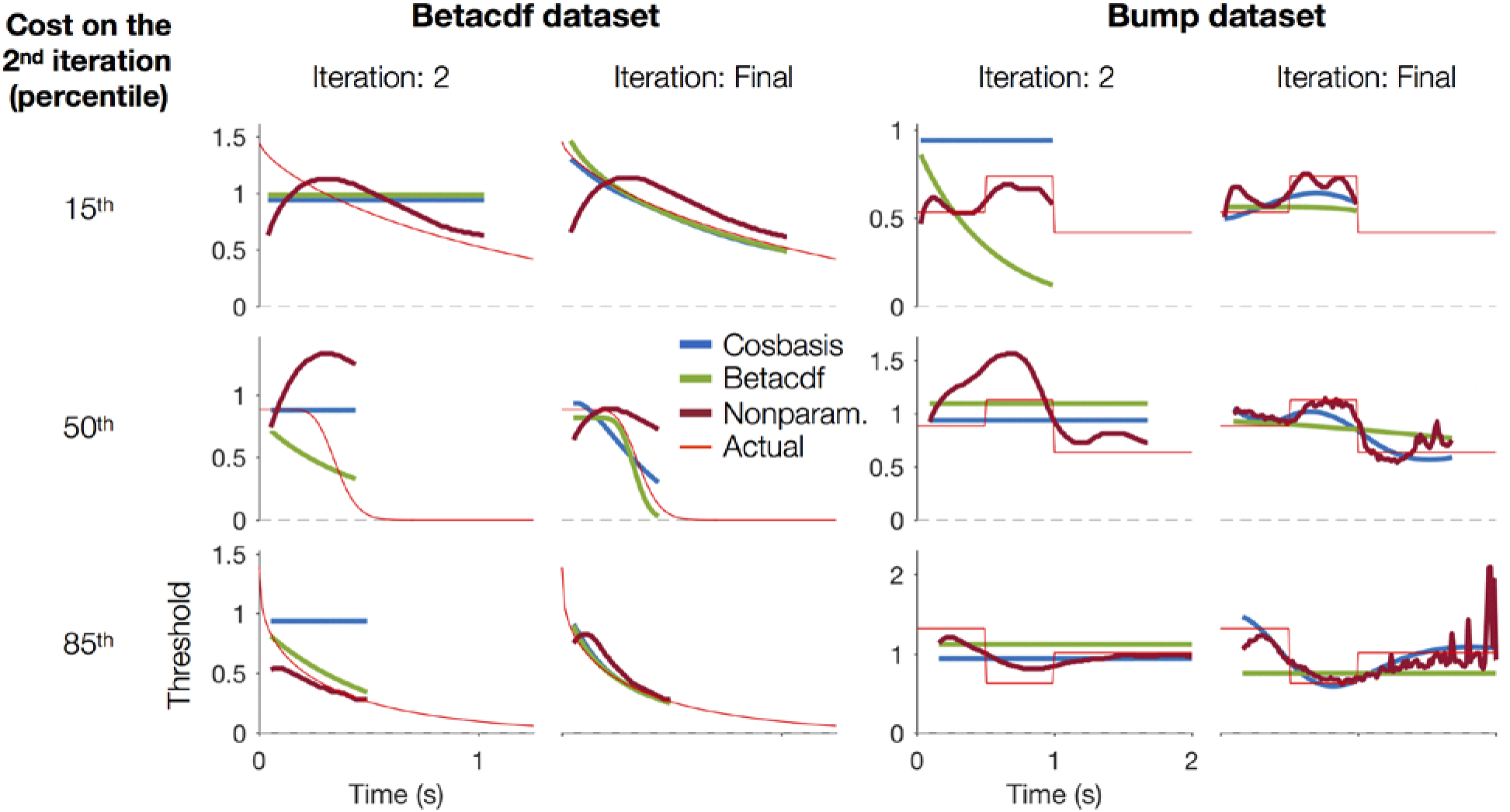
Example fits. *First column.* Example fits to the data simulated with the betacdf thresholds. The thin red line is the true threshold, and the thick blue, green, and red lines are the thresholds estimated by the cosbasis, betacdf, and nonparametric models on the 2^nd^ iteration of the gradient descent. The fit thresholds are drawn between 5^th^- and 95^th^-percentile decision times. The top row is the simulated dataset with the 15^th^-percentile cost for the nonparametric model on the 2^nd^ iteration of gradient descent, among all datasets. The second and the third rows are datasets with the 50^th^- and 85^th^- percentile costs. *Second column.* The fit thresholds after convergence, for the same datasets as the first column. *Third and fourth columns.* The same as the first two columns, except for the data simulated with the bump thresholds.

Finally, we compared the performance of the parametric and nonparametric fitting methods quantitatively. On the first column of Figure 6, we can see that all fitting methods have a high cost at the beginning of the fitting process, which decreases in later iterations. Notably, the nonparametric fits gave a lower average cost compared to parametric fits from the first iteration of the gradient descent for both betacdf and bump datasets (shuffle test: *p* ≤ 0.0002), and kept the lead for 11–15 iterations (Figure 6, first column; *p* < 0.05 up to iteration 11, 13, 15, and 11 compared to the combinations of the betacdf fit-betacdf dataset, cosbasis fit-betacdf dataset, betacdf fit-bump dataset, and cosbasis fit-bump dataset, respectively.)

**Figure 6.**
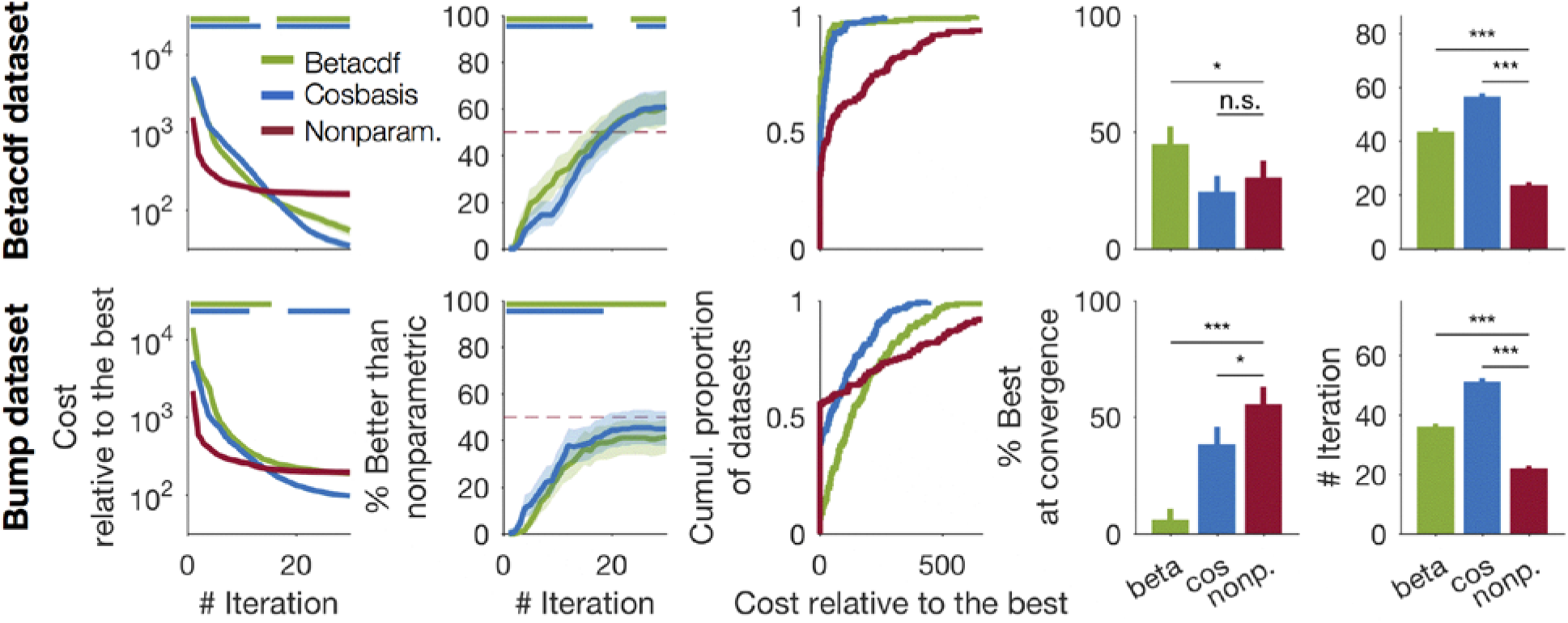
Comparison of performance between fitting methods. *First column.* Cost (relative to the smallest across all fitting methods after convergence within each simulated dataset) while fitting the betacdf and bump datasets, as a function of the number of iterations (see Methods). Lines and shades are the mean and SEM (which was often smaller than the thickness of the lines). Green, blue, and red correspond to the betacdf, cosbasis, and nonparametric fits, respectively. Horizontal lines at the top indicate the intervals where the difference is significant (*p* < 0.05, two-tailed), determined by 10,000 random pairwise shuffling of the fitting method label with the nonparametric method. *Second column.* Percentage of the simulated dataset for which the given parametric method gave lower cost than the nonparametric method. Lines and shades are the mean and the 95% binomial CI. The dashed horizontal line marks the 50% level. Horizontal lines at the top indicate the intervals where the difference from 50% is significant (*p* < 0.05, two-tailed), determined as above. *Third column.* Cumulative distribution of the cost relative to the best (see above) after convergence. Rightmost parts of the graphs are truncated. *Fourth column.* Proportion of the datasets for which each fitting method gives the lowest cost (across the 3 methods) at convergence (%) and the 95% CI. Markers indicate the significance of pairwise comparisons (sign test; ***: *p* < 0.001; **: *p* < 0.01; *: *p* < 0.05; n.s.: p > 0.1). *Fifth column.* Average number of iterations needed for convergence (± SEM). The meaning of the markers is the same as above.

We also evaluated the nonparametric fits in terms of the *proportion* of the simulated datasets (each with a random parameter setting) for which the nonparametric fits gave a higher cost than the parametric fits on each iteration (Figure 6, second column). The results favored the nonparametric fits more than the average cost did: the nonparametric fits gave a better fit in significantly more datasets than the parametric fits on the first iterations (shuffle test: *p* < 0.05 up to the first 15, 16, 133, and 18 iterations compared to the combinations of the betacdf fit-betacdf dataset, cosbasis fit-betacdf dataset, betacdf fit-bump dataset, and cosbasis fit-bump dataset, respectively.) By either metric (average cost or proportion of lower cost), the nonparametric model was a significantly better choice than the parametric models if we were to restrict the number of iterations allowed for fitting.

Not surprisingly, for the betacdf dataset, the nonparametric fits were not better than the parametric fits at convergence. Nonparametric fits gave the smallest cost less frequently than the betacdf fits (Figure 6, fourth column, top row; *p* = 0.03), although it did not differ significantly from cosbasis fits (*p* = 0.31), and its average cost was higher (paired t-test: *p* < 0.001) compared to both betacdf and cosbasis fits. This is expected since the parametric models are constrained to have smoothly-varying thresholds, as is the model that generated the dataset.

However, for the bump dataset, the nonparametric fits gave the best fit significantly more frequently than either the betacdf or the cosbasis fits (Figure 6, fourth column, bottom row; sign test: *p* < 0.001 and *p* = 0.02), although it converged significantly faster (Figure 6, rightmost column, bottom row; paired t-test: *p* < 0.001 compared to both betacdf and cosbasis fits).

Its advantage in the speed was qualitatively the same (the order among fitting methods was the same and the p-values were all < 0.001), when we tried comparing (1) the total execution time, or (2) the “pure” execution time devoted to the calculation of cost, by multiplying the number of iteration with the time per calculation of cost. Here, the time per calculation of cost were 67 ± 8, 70 ± 4, and 91 ± 5 ms [mean ± SD] for the betacdf, cosbasis, and nonparametric models, based on 100 calculations after 3 warm-up calculations. The ratio of the time compared to the nonparametric model’s were 74% and 77% for the betacdf and cosbasis models.

Interestingly, the average cost of the nonparametric fits at convergence for the bump dataset was higher than that of the cosbasis fits (paired t-test: *p* < 0.001), although it was not significantly different from that of the betacdf fits (*p* = 0.57). Inspection of the cumulative distribution function of the cost at convergence showed why (Figure 6, third column, bottom row). The nonparametric fits gave the lowest cost most frequently (red), but when it was not the best, it tended to give higher cost than the parametric models. We discuss the implication of the characteristics of the nonparametric model’s performance below.

## Discussion

We showed that we can estimate time-varying decision thresholds without an assumption on their shape with a simple heuristic algorithm. The method does not require any hyperparameter for the threshold, yet it showed significantly better performance compared to parametric methods when a limited number of gradient descent steps are run: during the initial steps, it was even better than the parametric models of the same functional form as the one generated the data (Figure 6, first column). At convergence, the method performed better than parametric methods more frequently when the shape of the thresholds of the model that generated the data did not match the parametric models’ (Figure 6, fourth column, bottom row), although it needed fewer iterations to converge (Figure 6, rightmost column, bottom row).

Given the method’s initial advantage in performance from the first step of the gradient descent, it would be especially useful for situations where computing time is extremely limited, such as for online analyses of the data from high-throughput neural recording, financial markets, or large-scale internet-based services. Notably, the new method showed the initial advantage even compared to models whose functional form of the threshold matched what generated the data. If this comparative advantage holds for other functional forms as well, we may be able to choose the new method without worrying about exactly what kind of threshold generated the data, as long as we need to limit the fitting time enough.

When the shape of the true thresholds did not match the shape assumed by the parametric models, the nonparametric model performed the best on convergence more frequently than the parametric models, but when its fit was bad, the cost tended to be higher (Figure 6, third column, bottom row). Such performance may be suited for applications to a competitive situation where being the best model is the only important consideration (i.e., when it does not matter how bad the fit is when it is not the best among all.) To improve the model’s fits in the future, it may be worthwhile to examine if there is a particular regime of the parameter settings that the nonparametric model fails to fit.

In this preliminary report, we did not use cross-validated costs, and the obvious next step for us is to confirm the results with cross-validation. However, we suspect that the results shown would not change qualitatively, because our conclusions are based on simulated data where we know the true thresholds, and the match (or mismatch) of the fitted thresholds (Figure 5) with the true thresholds is unlikely to have been driven entirely by outliers.

There are a number of directions that the method can be extended. The simplest would be to introduce a smoothness prior. The appropriate smoothness may be found using cross validation. Another direction is to use a better method to estimate the decision time distribution from the reaction time distribution than the current one (Eq. 20) which is approximate and biased (see Appendix). While many other directions are possible, it would need careful consideration to retain the advantage in speed, because if the speed is not a concern we can simply resort to parametric models (e.g., the cosbasis model with many bases). In this regard, it may be worthwhile to look for a way to get the best of both worlds. For example, the method may be combined with parametric models, by quickly providing information about the functional form. We may first run the nonparametric model for a few iterations to find out around how much time after the stimulus onset the threshold changes the most, and subsequently fit the data with cosine basis functions, with more basis functions placed around the time of the most change. Such elaboration may improve the fit while preserving the speed. The method we described here may serve as a simple yet useful starting point for more elaborate methods.

## Appendix Estimating the decision time distribution from reaction times and θ_N_

In Eq. 19, we estimate the distribution of the decision time *T*_D_ from the observed reaction times (included in the data **D**) and parameters for the nondecision time **θ**_N_ for the case with choice *z*, which is reproduced here:

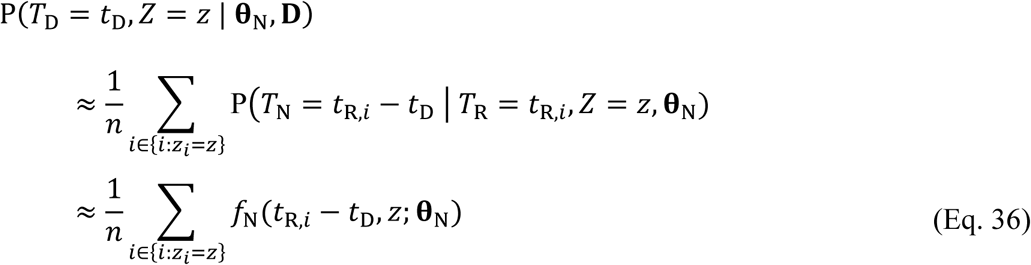

To simplify the problem without losing generality, let’s condition all variables on *Z* = *z*:

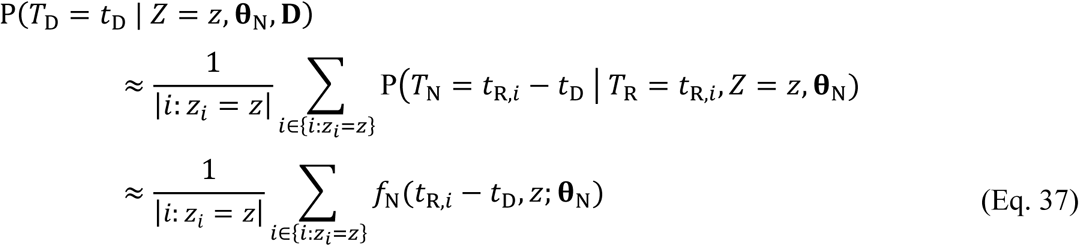

Here, |⋅| denotes the cardinality, division with which ensures that the right-hand side sums to 1. The decision time distribution computed this way is not guaranteed to maximize the probability of observing the data given **θ**_N_. To see this, consider Eq. 1, reproduced here with asterisks attached, to denote that the symbols relate to the true random variables of the generative model, in the case where the choice is *z*:

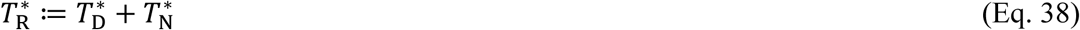

Let’s assume that our **θ**_N_ matched the generative model’s. Then, if we denote with 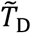 the *T*_D_ we calculated above with Eq. 37, and with 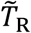 the empirical distribution {*t*_*R,i*_} in the case *i* ∈ {*i*: *z*_*i*_ = *z*}, then Eq. 37 can be rewritten as:

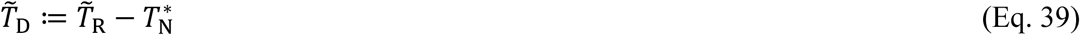

This way of calculation is nice in that the expectation of 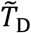 matches the true value:

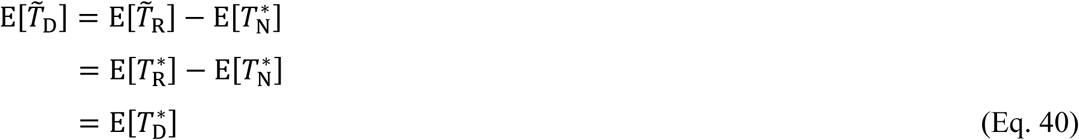

However, the expectation of the variance 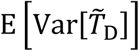 is bigger than the true value, since variance is added even when the random variable is subtracted, and since we are interested in the case where 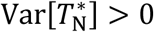:

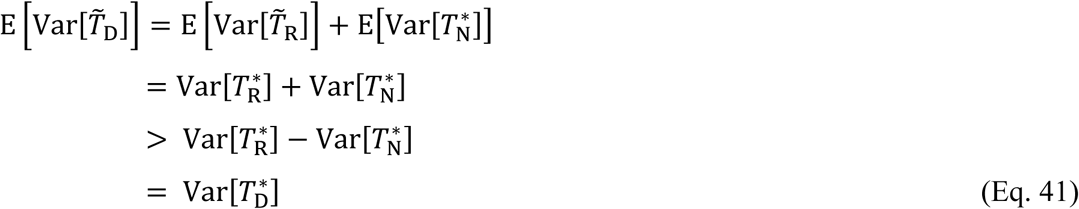

The last equality comes from applying Var[⋅] to both sides of Eq. 38 and rearranging. Therefore, we expect 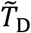 to be biased to have too big a variance.

There may be better ways to estimate *T*_D_ given **θ**_N_ and **D**. We can imagine using some kind of deconvolution of 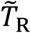, probably after smoothing for numerical stability. We can also imagine parameterizing the distribution of *T*_D_ with, e.g., cosine basis functions, and optimizing it to give the best likelihood of observing the data, through steps like Eqs. 2–3. However, we chose to use Eq. 36 (i.e., Eq. 19) due to its simplicity and speed. There certainly could be a better method, and we defer finding it to future works. We nonetheless confirmed that Eq. 36 gives a better-than chance approximation of the true distribution of 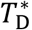, as follows.

To see if Eq. 36 (i.e., Eq. 19) gives a better-than-chance approximation of the true simulated decision time distribution, we computed the Kullback-Leibler divergence:

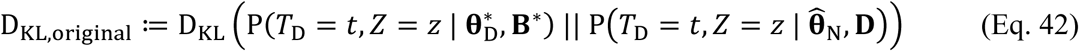

Here, 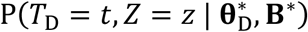 is the distribution from the generative model, computed with Eqs. 14–15 with the **θ**_D_ and **B** used in simulating the data (denoted by asterisks), as described in Section “Comparison of the performance with the parametric models”. 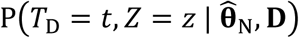 is computed as in Eq. 36 with the value of **θ**_N_ estimated by the new (nonparametric) method at convergence, 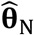. We computed D_KL,original_ for all 180 “bump” datasets, then examined their difference from D_KL,shuffled_, which are computed after shuffling the association between 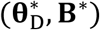 and 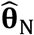 across the 180 datasets. The results clearly showed that D_KL,shuffle_ tends to be bigger than D_KL,original_ (sign test; *p* < 10^−15^; Figure 7), confirming that Eq. 36 (i.e., Eq. 19) gives a better approximation of the original decision time distribution than of a randomly chosen decision time distribution. The performance is definitely not perfect (only 89% of D_KL,original_ was better than D_KL,shuffle_): it may be due to similarity between the decision time distributions, bias in the algorithm as described above, or likely, both. We defer it to future works to improve the estimation of the decision time distribution.

**Figure 7.**
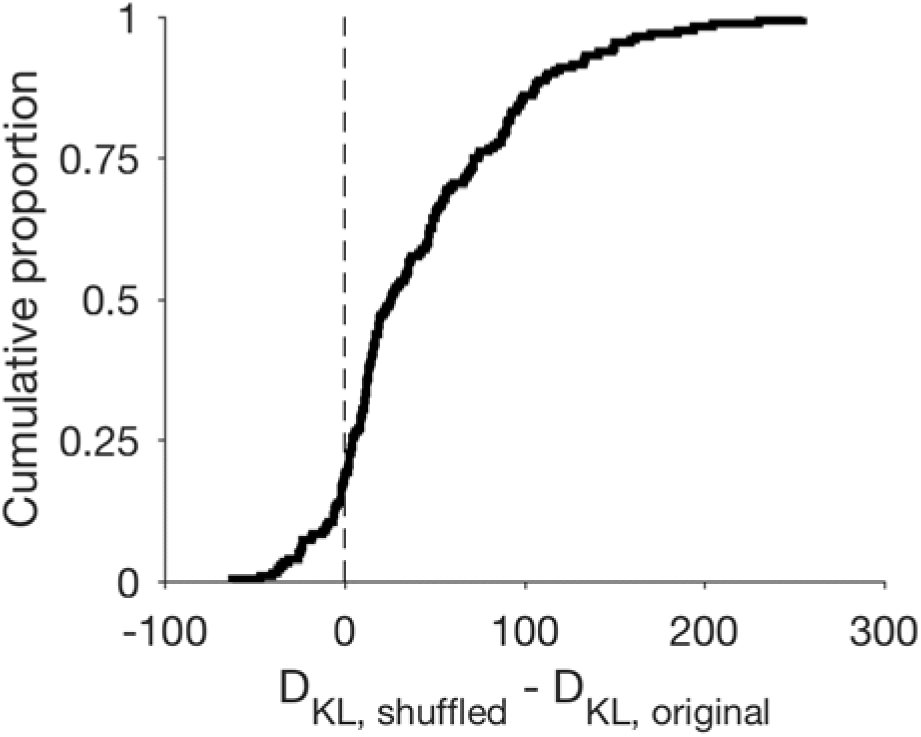
Cumulative distribution of the difference in D_KL_. See above text and Eq. 42. Positive value on the abscissa indicates that the original is better.

